# Synthesis-free PET imaging of brown adipose tissue and TSPO via Combination of Disulfiram and ^64^CuCl_2_

**DOI:** 10.1101/131193

**Authors:** Jing Yang, Jian Yang, Lu Wang, Anna Moore, Steven H. Liang, Chongzhao Ran

## Abstract

PET imaging is a widely applicable but a very expensive technology. Strategies that can significantly reduce the high cost of PET imaging are highly desirable both for research and commercialization. On-site synthesis is one important contributor to the high cost. In this report, we demonstrated the feasibility of a synthesis-free method for PET imaging of brown adipose tissue (BAT) and translocator protein 18kDa (TSPO) via a combination of Disulfiram, an FDA approved drug for alcoholism, and ^64^CuCl_2_ (termed ^64^Cu-Dis). Our blocking studies, Western blot, and tissue histological imaging suggested that the observed BAT contrast was due to ^64^Cu-Dis binding to TSPO, which was further confirmed as a specific biomarker for BAT imaging using [18F]-F-DPA, a TSPO-specific PET tracer. Our studies, for the first time, demonstrated that TSPO could serve as a potential imaging biomarker for BAT. Furthermore, since imaging contrast obtained with both ^64^Cu-Dis and [18F]-F-DPA was not dependent on BAT activation, these agents could be used for reliably imaging BAT mass. Additional value of our synthesis-free approach could be applied to imaging TSPO in other tissues as it is an established biomarker of neuro-inflammation in activated microglia and plays a role in immune response, steroid synthesis, and apoptosis. Although here we applied ^64^Cu-Dis for a synthesis-free PET imaging of BAT, we believe that our strategy could be extended to other targets while significantly reducing the cost of PET imaging.

**Significance:** Brown adipose tissue (BAT) has been considered as “good fat,” and large-scale analysis has undoubtedly validated its clinical significance. BAT tightly correlates with body-mass index (BMI), suggesting that BAT bears clear significance for metabolic disorders such as obesity and diabetes. BAT imaging with [18F]-FDG, the most used method for visualizing BAT, primarily reflects BAT activation, but not BAT mass. A convenient imaging method that can consistently reflect BAT mass is still lacking. In this report, we demonstrated that BAT mass can be reliably imaged with a synthesis-free method using the combination of Disulfiram and ^64^CuCl_2_ (^64^Cu-Dis) via TSPO binding. We further demonstrated for the first time that TSPO is a specific imaging biomarker for BAT.

## Introduction

Positron emission tomography (PET) has been widely used for clinical and preclinical studies, including disease diagnosis, treatment monitoring, and drug development. Compared to other imaging modalities, such as MRI, ultrasound, and optical imaging, PET is highly sensitive and quantitative (1). However, the high cost of PET imaging has been the key roadblock for its widespread routine use. The high cost originates from expensive radionuclide production and tracer preparation, which must be conducted on-site (1). Here, we demonstrated the feasibility of a synthesis-free PET imaging method for brown adipose tissue (BAT). With this method, the cost of PET imaging could be dramatically reduced, thus allowing for a widespread application of this technology.

For the purpose of synthesis-free PET imaging, we considered the following criteria: 1) the radionuclide should have a suitable lifetime for delivery and transportation; 2) synthetic components (excluding the radionuclide) should be available in a convenient kit; and 3) the generation of the actual tracers should be very fast or the actual tracers should be formed in vivo. To meet the above requirements, we selected ^64^Cu as the radionuclide due to its reasonable decay lifetime (12.7 hours) and widespread availability. For our proof-of-principle studies we selected BAT as the biological target, because BAT has the following unique features making it a suitable imaging model (2, 3): 1) BAT in mice is situated away from large organs such as liver, heart, and stomach, and thus signal interference from these large organs is minimal; 2) BAT is a whole mass organ; 3) BAT has a unique triangular physical shape which is easy to distinguish from other tissues.

Besides the above features, clinical significance of BAT has been validated through a large-scale analysis of [18F]-FDG PET-CT images and other important studies (4-8). BAT is a specialized tissue for thermogenesis in mammals, whose function is to dissipate large amounts of chemical/food energy as heat, thus maintaining the energy balance of the whole body (9-11). In spite of the fact that investigations of BAT have been ongoing for 70 years, it had been assumed that BAT disappears from the body of adults and has no significant physiological relevance in adult humans (9, 12-14). This “non-existence” assumption is partially due to the lack of proper imaging methods to “see” the small BAT depots *in vivo*, as only 3%-8% of BAT depots in adults could be clearly visualized with [18F]-FDG (the most used imaging method) if no cold or drug stimulation is applied (6, 15-17). However, under stimulated conditions, [18F]-FDG PET imaging has shown that BAT is still present in 95% of healthy adults in the upper chest, neck, and other locations (4-6). This remarkably large difference between unstimulated and stimulated conditions strongly indicates that [18F]-FDG PET imaging primarily reflects the activation of BAT, but not BAT mass. Various other imaging methods for BAT are available for preclinical and clinical studies, however most of them are dependent on BAT activation (18-27). Therefore, an imaging probe that can consistently report on BAT mass is highly desirable.

In this report, we first conducted a top-down screening, and found that the combination of <u>^64^Cu</u>Cl_2_ and <u>Dis</u>ulfiram (termed ^64^Cu-Dis) provided high contrast for BAT, which was not affected by BAT activation. We also found that TSPO, a transport protein located on the outer mitochondrial membrane (28), was the binding target of ^64^Cu-Dis. We further validated that TSPO is an excellent but unexpected imaging biomarker for BAT using Western blot, histology, and PET imaging with TSPO-specific ligand [18F]-F-DPA.

## Results

### 1. Screening for BAT-binding ligands

In our previous report (3), we have demonstrated that a top-down screening approach could be used for seeking near infrared fluorescence (NIRF) imaging probes for BAT. In that study, we screened 38 NIRF dyes resulting in two hits that we further optimized for high BAT selectivity (3). In the present report, we used a similar top-down strategy for fast screening of a library of copper ligands, which could be used for fast coordination chemistry with no need for purification. Among the 16 screened ligands, four compounds were considered as positive hits (SI Fig.1), including Disulfiram, diethyldithiocarbamate (DDC, a metabolic product of Disulfiram in vivo (29)), cysteamine, and salicylaldoxime.

**Fig. 1.**
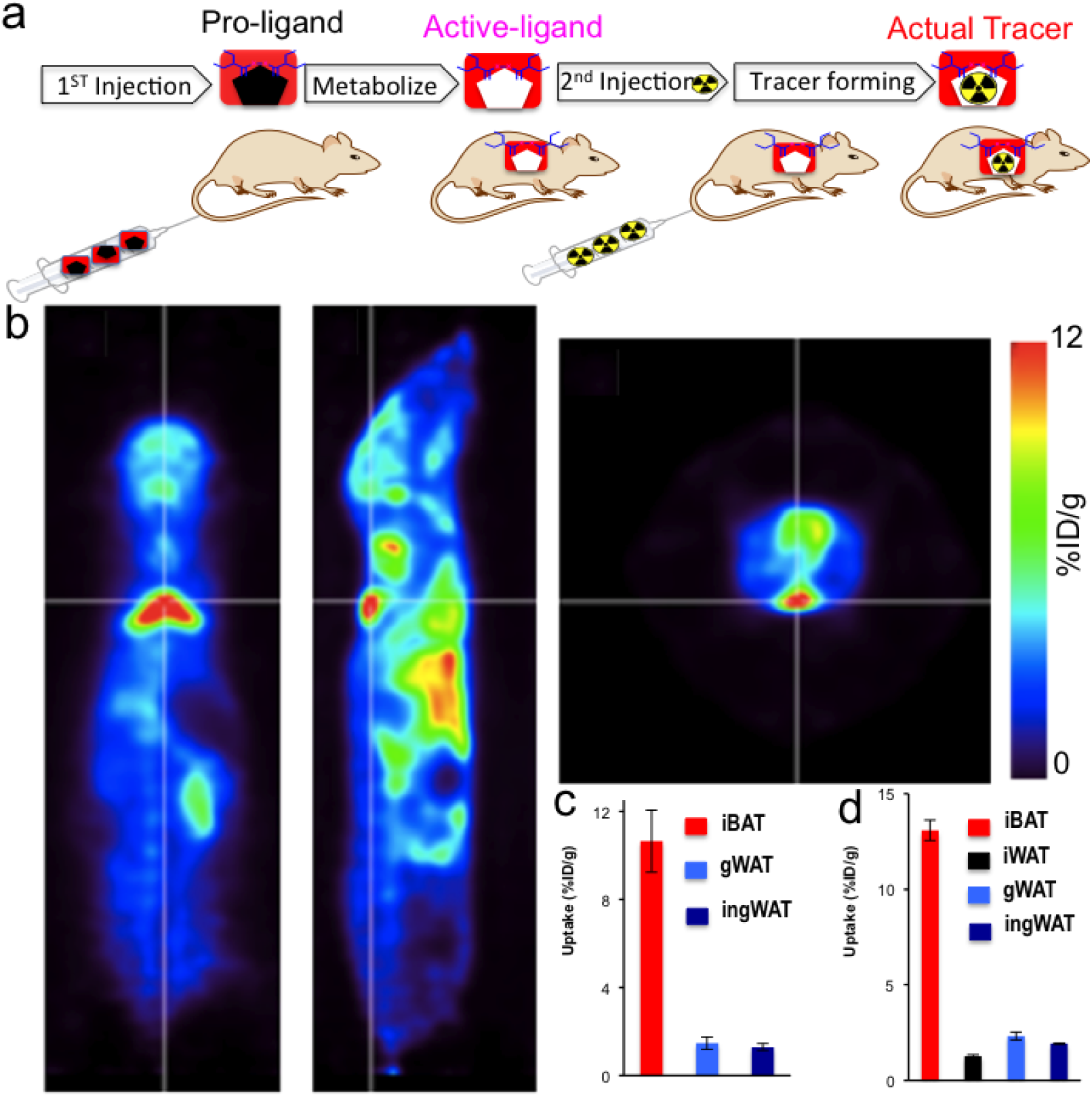
(a) Diagram of synthesis-free PET imaging via stepwise injection. The pro-ligand is first injected to metabolize in vivo to produce the active ligand, which will bind the lately injected radionuclide to form actual PET tracer in situ. (b) Representative PET images of a mouse injected consecutively with Disulfiram and ^64^CuCl_2_ (^64^Cu-Dis) 30 minutes after ^64^CuCl_2_ injection. Coronal view (left), sagittal view (right), and transverse view (upper corner). The triangular shape of BAT can be easily identified. (c) Quantitative analysis of the tracer uptake from in vivo imaging with ^64^Cu-Dis in interscapular BAT (iBAT), gonadal WAT (gWAT) and inguinal WAT (ingWAT). (d) Ex vivo bio-distribution of the tracer in iBAT, interscapular WAT (iWAT), gWAT and ingWAT.

### 2. Using the combination of ^64^CuCl_2_ and Disulfiram (^64^Cu-Dis) for “synthesis-free” PET imaging

One of the lead ligands, Disulfiram, caught our attention, because it is an FDA approved drug for alcoholism (30). Disulfiram, a disulfide compound, does not have a strong affinity to copper. However, when injected in vivo, it is reduced to the monomer and releases the thiol group producing diethyldithiocarbamate (DDC), which has a high affinity for copper (II) (31). Considering that DDC, a metabolic product of Disulfiram, is an active copper chelator, we proposed a step-wise injection strategy to realize synthesis-free PET imaging (Fig.1a). To this end, we injected Disulfiram intraperitoneally (40 mg/kg) in mice and allowed 60 minutes for it to metabolize into DDC, the active copper binding form. Next, we intravenously injected ^64^CuCl_2_, which can quickly chelate with DDC in vivo. Strikingly, we found that BAT could be easily identified by PET imaging as early as 10 minutes (Fig.1b), and as late as 48 hours after injections. The uptake of the tracer at 30 minutes after ^64^CuCl_2_ injection was 10.6 %ID/g (Fig.1c). More importantly, no apparent signal could be observed from WAT (white adipose tissue), including inguinal and epidermal areas (Fig.1c and SI Fig.2), indicating that this method has high BAT selectivity over WAT.

Attaining high BAT/WAT selectivity has been one of the challenges for developing imaging probes for BAT. To confirm the data on BAT/WAT selectivity obtained from in vivo PET imaging, we conducted ex vivo bio-distribution of BAT and WAT tissues 6 hours after ^64^CuCl_2_ injection. We found that interscapular BAT uptake was 13.1 %ID/g, which was 10-fold higher than that of interscapular WAT, inguinal WAT, or gonadal WAT. (Fig.1d). Taken together, our in vivo imaging data were consistent with the ex vivo data, suggesting that ^64^Cu-Dis was highly selective for BAT over WAT.

We also conducted control experiment with ^64^CuCl_2_ only, and found that BAT uptake was about 1.0% ID/g (SI Fig.3), which was much lower than that with ^64^Cu-Dis, suggesting that Disulfiram is necessary for high BAT uptake.

### 3. In vivo BAT imaging with ^64^Cu-Dis

To investigate whether the “synthesis-free” method can be used to consistently image BAT mass under different conditions, we imaged mice under a normal condition and under cold exposure, which is a standard protocol for BAT activation (2, 32). For cold exposure, the mice were placed in a cold room at 4°C for 2 hours before and 1 hour after Disulfiram administration. Next, the mice were injected with ^64^CuCl_2_ and kept at 4°C for 30 minutes followed by imaging for 30 minutes (Fig.2a). BAT uptake was about 10.0 % ID/g for normal conditions at 1 hour after ^64^CuCl_2_ injection with no significant decrease at 6 hours (SI Fig.4). Importantly, we found that there was no significant difference in uptake between normal and cold treated groups (Fig.2b,c). This is contrary to [18F]-FDG PET and other imaging methods, in which BAT contrast is highly dependent on its activation status. Our data suggested that this synthesis-free method could be used for reporting BAT mass regardless of BAT activation status.

**Fig. 2.**
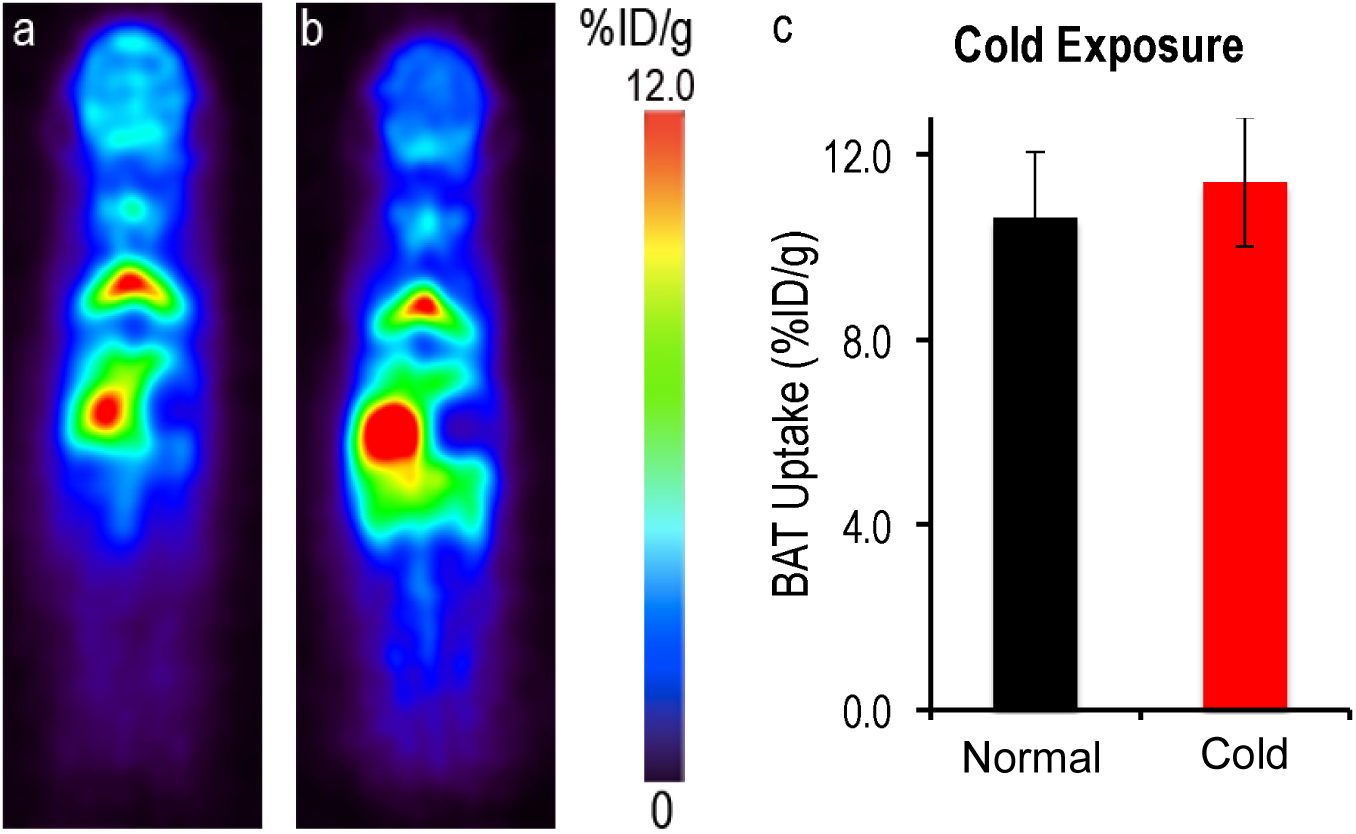
PET images of mice injected ^64^Cu-Dis under normal and cold exposure conditions. (a) Representative coronal image under a normal condition, (b) Representative coronal image under a cold exposure condition, and (c) Quantitative analysis of (a) and (b). There was no significant difference in uptake between control and cold treated groups (p = 0.359).

It has been reported that different anesthesia regimens have significant suppression impact on BAT activity (7, 33, 34). To investigate this, we kept mice under light anesthesia for 20 minutes before and 40 minutes after Disulfiram injection, followed by ^64^CuCl_2_ injection. We found that there was no significant difference in BAT uptake in animals under 20-minute anesthesia compared to animals under normal conditions (SI Fig.5), again suggesting that BAT imaging with ^64^Cu-Dis is not dependent on BAT activity.

### 4. Blocking studies with TSPO-specific ligand F-DPA

Our in vivo and ex vivo results revealed that the ^64^Cu-Dis combination was indeed a synthesis-free method for PET imaging of BAT, however, it was not clear what the molecular binding target was. By surveying the publications reporting on the Disulfiram mechanism of action, and found that TSPO could be a target candidate (35, 36). To investigate whether the uptake of Disulfiram or DDC is related to TSPO, we conducted blocking studies using the TSPO specific ligand F-DPA (Ki = 9nM) (Compound 3K in reference 37, and structure in SI Fig.6c) (37). F-DPA is an analogue of [F18]-DPA-714 (38), and is a highly selective ligand for TSPO (37, 39). In these experiments, we intravenously injected 3.0 mg/kg of F-DPA after ip injection of Disulfiram, and imaged the mice using the same protocol as above. Images were captured at 1 hour and 6 hours after ^64^CuCl_2_ injection. Indeed, we found that F-DPA could effectively block BAT signal at both time points (Fig.3a,b), with a 35.0% and 60.0% decrease respectively (Fig.3c,d). Interestingly, we also observed a significant decrease in PET signal from other organs such as heart and kidney (Fig.3e,f and SI Fig.6a,b), in which TSPO has reportedly high expression (40). Therefore, our data strongly suggested that high BAT contrast was associated with TSPO.

**Fig. 3.**
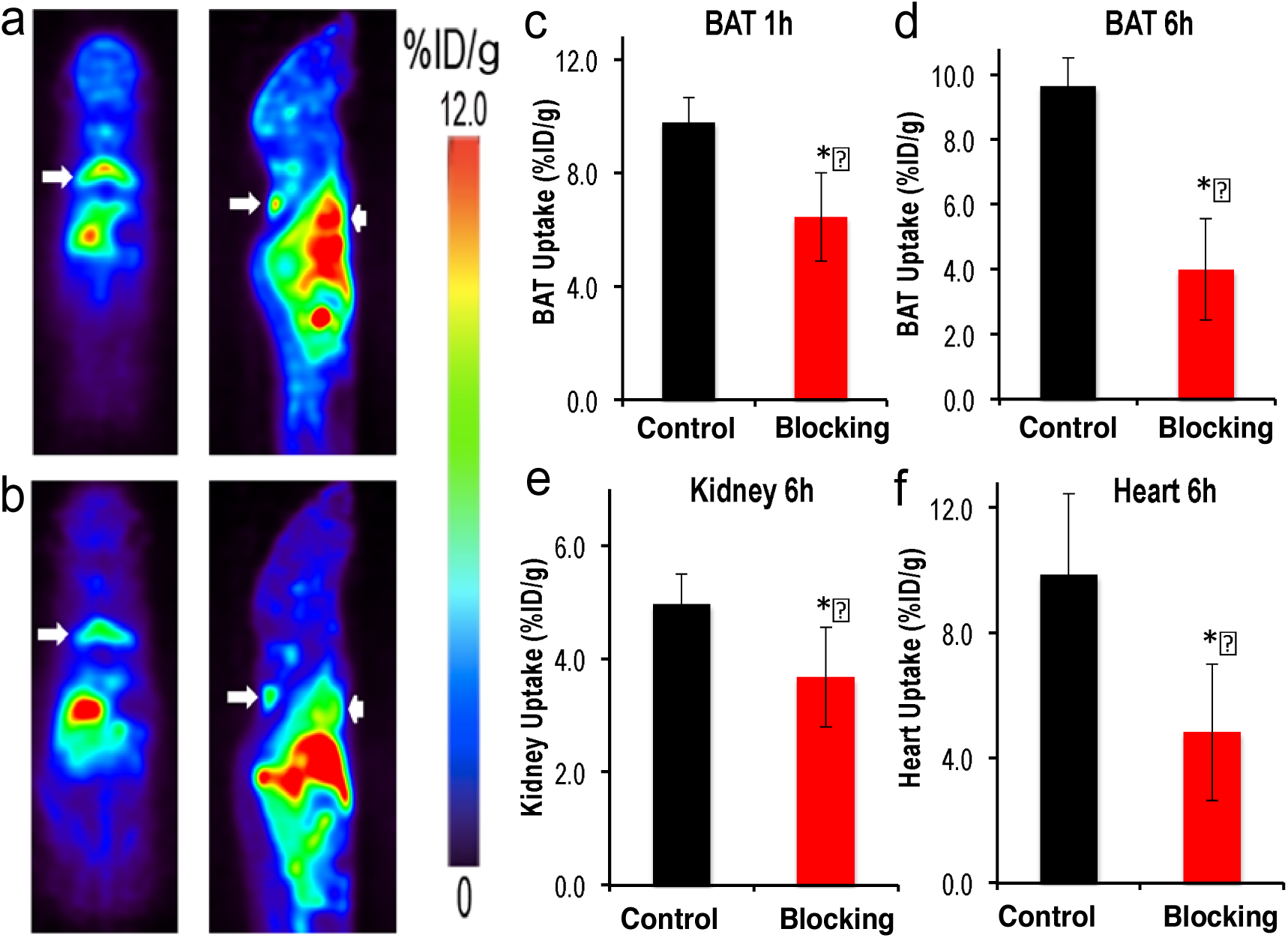
Blocking studies with TSPO ligand F-DPA. (a) Representative PET images of mice injected with ^64^Cu-Dis without blocking. (b) Representative PET images of mice with ^64^Cu-Dis with F-DPA blocking. The decreased uptake of ^64^Cu-Dis can be observed in BAT (white arrow) and heart (arrow head). (c) Quantitative analysis of BAT uptake at 1 hour, (d) Quantitative analysis of BAT uptake at 6 hours, (e) Quantitative analysis of kidney uptake at 6 hours, and (f) quantitative analysis of heart uptake at 6 hours. * indicates p < 0.05.

### 5. Biological validation of TSPO as a specific biomarker for BAT imaging

To further confirm that TSPO can be used as a specific imaging biomarker for BAT, we performed biological analyses. TSPO, also called translocator protein or peripheral benzodiazepine receptor (PBR) (28, 40), is located on the outer mitochondrial membrane, and one of its characteristic features of BAT is its high abundance of mitochondria compared to WAT (9-11). It is likely that the abundance of TSPO in BAT is much higher than in WAT. Nonetheless, no experimental data are available indicating whether TSPO expression in BAT and WAT is significantly different, which is crucial for determining whether TSPO can be used as a specific imaging biomarker for BAT. To answer this question, we compared the qPCR data of TSPO in BAT and WAT, and found that the mRNA level in BAT was about 1.5-fold higher than in WAT (SI Fig.7). To further confirm different levels of TSPO protein expression in BAT and WAT, we performed Western blot with anti-TSPO antibody. As shown in Fig. 4a, protein level in BAT was about 20-fold higher than in WAT (Fig.4a,b). We next investigated the abundance of mitochondria in BAT and WAT tissues using a mitochondria specific dye (MitoTrack deep red FM). As expected, BAT showed much higher fluorescence intensity than WAT (Fig.4c). In addition, we performed immunological staining of BAT and WAT tissues with anti-TSPO antibody. Fig. 4d demonstrated that the abundance of TSPO in BAT was significantly higher than that in WAT. In conclusion, our data for the first time suggest that TSPO could be used as a specific imaging biomarker for BAT.

**Fig. 4.**
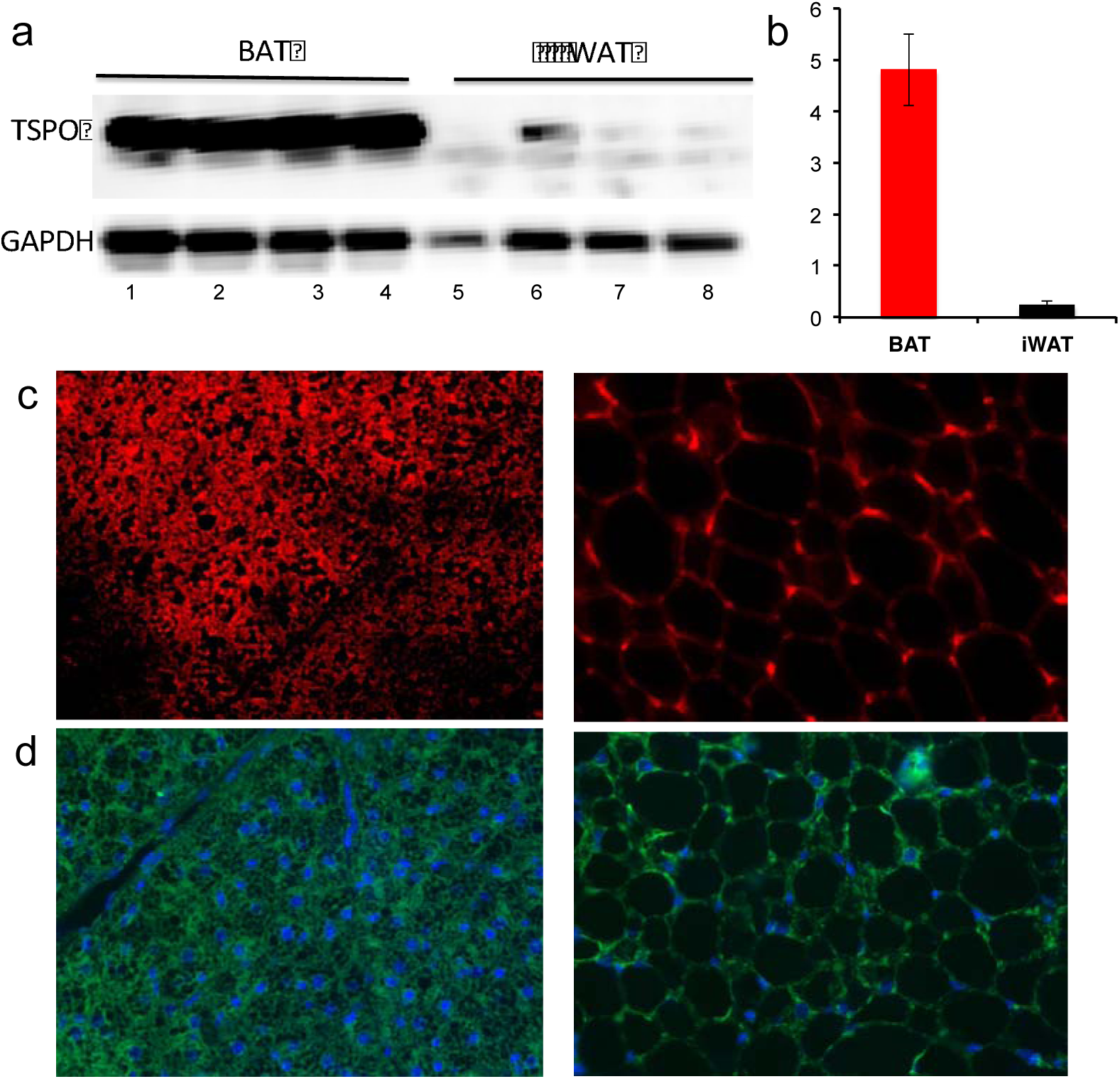
Biological validation of high expression level of TSPO in BAT. a) Western blot of BAT and WAT tissue extractions with anti-TSPO antibody (lane1-4: four duplicated BAT samples; lane5-8: four duplicated WAT samples), b) quantification of a). Note higher TSPO level in BAT compared to WAT. c) Histological staining of BAT (left) and WAT (right) tissue slices with mitochondria specific dye MitoTrack deep Red, and d) immunohistological staining of BAT and WAT tissue with anti-TSPO antibody (green) and DAPI for nuclei (blue).

### 6. PET imaging of BAT with TSPO-specific tracer [18F]-F-DPA

To further validate whether TSPO is a specific imaging biomarker for BAT, we investigated whether existing TSPO-specific PET tracers could provide high contrast for BAT over WAT. To this end, we used 18F-labeled TSPO-specific ligand [18F]-F-DPA (37-39) We first performed PET imaging in Balb/c mice using a 20-minute static scan. As seen in Fig. 5a-c, there was a readily identifiable contrast from BAT with a 16% ID/g uptake 30 minutes after the injection. Moreover, we also compared images at locations with high WAT abundance and found that inguinal and gonadal WAT had minimal uptake (SI Fig.8).

**Fig. 5.**
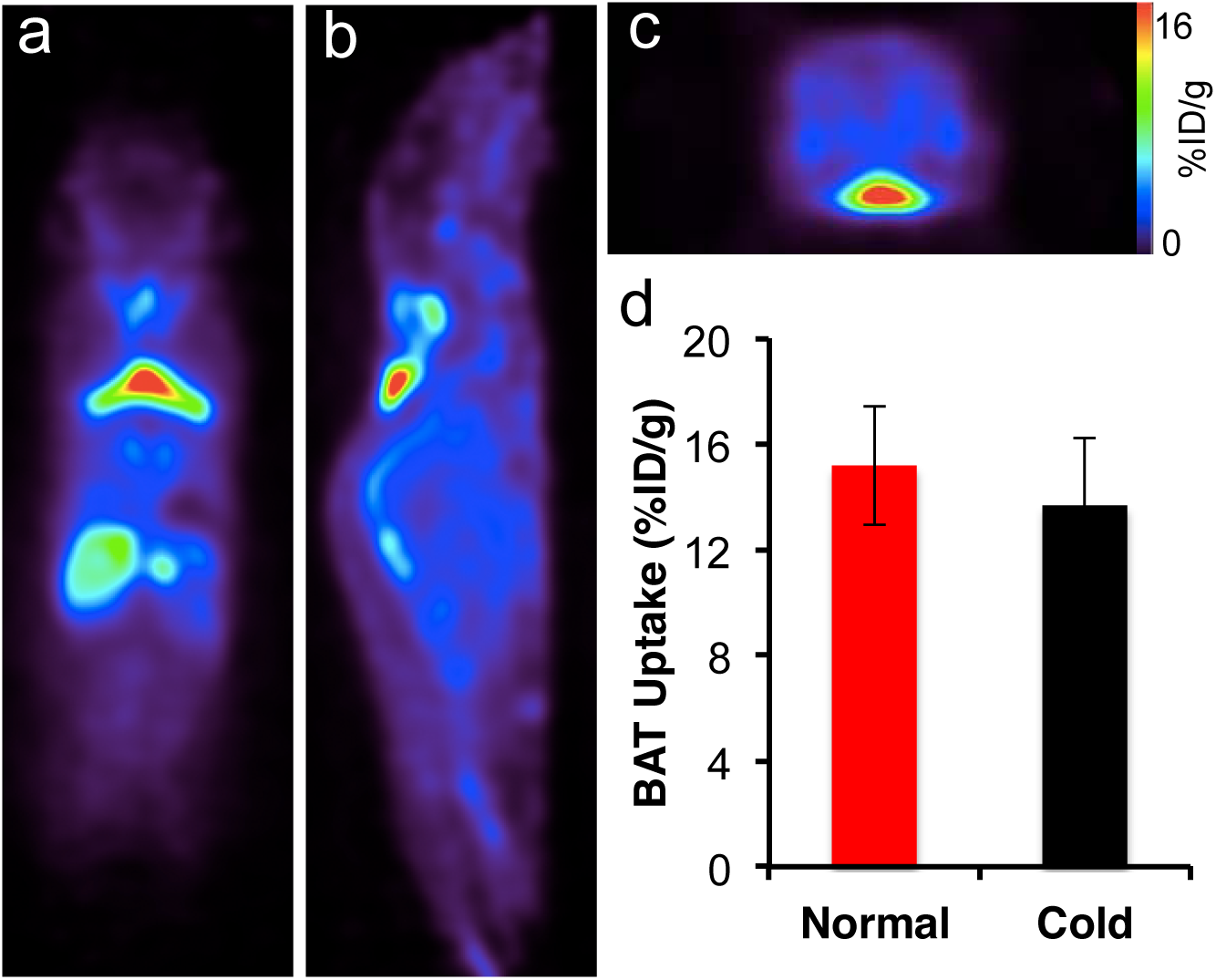
Representative PET images of mice injected with 18F-DPA. (a) Coronal view, (b) sagittal view, and (c) transverse view. The triangular shape of BAT can be clearly identified. (d) Quantitative analysis of BAT uptake under normal and cold exposure. There was no significant difference between these two groups (p = 0.356).

To further validate the uptake of [18F]-F-DPA in BAT, we conducted a bio-distribution study in Balb/c mice at 45 minutes after the probe injection. BAT, inguinal WAT and gonadal WAT, brain, liver, and heart were harvested (SI Fig.9). The bio-distribution data confirmed a higher uptake of the tracer in BAT compared to other organs, suggesting that [18F]-F-DPA was highly selective for BAT compared to WAT. Taken together, the high uptake in BAT with established TSPO PET ligand supports our hypothesis that TSPO could serve as an imaging biomarker for BAT.

### 7. 18F-DPA uptake is not influenced by BAT activation

To investigate whether [18F]-F-DPA can be a reliable PET tracer for BAT mass, we conducted PET imaging under BAT activation with cold exposure. Mice were exposed to 4°C for 2 hours before imaging. Our imaging data showed that there were no significant differences between cold treatment and normal condition, indicating that [18F]-F-DPA could serve as a reliable tracer for BAT mass independent of its activation (Fig.5d). Thompson et al have recently showed that TPSO expression could not be altered by acute cold exposure, which is consistent with the data obtained in our experiment (41). This also suggests that TSPO is a reliable biomarker for BAT mass.

## Discussions and conclusions

PET imaging is immensely useful for both clinical and preclinical applications, however its high cost prevents it from being widely applicable. Requirements for an on-site cyclotron and on-site synthesis are among the major contributors of the high cost. In this report, we demonstrated the feasibility of a synthesis-free PET imaging. Although we demonstrated the application of this method for a specific case that involved imaging of BAT mass, we believe that “pseudo-synthesis-free” and “synthesis-free” methods could be used for other applications as well. “Pseudo-synthesis-free” methods can be realized with fast and strong chelation reaction of ligands with suitable radionuclides. “Synthesis-free” could be feasible if some compounds, which have strong coordination with the radionuclides, could pre-target receptors/enzymes, or if peptides/proteins that have strong chelation with radionuclides could be engineered into specific targets.

In the present study, Disulfiram was used as a precursor of a copper ligand, and the actual PET tracer formed in vivo via the chelation of ^64^CuCl_2_ with DDC, the in vivo metabolite of Disulfiram. In 2014, Nomura et al reported that Disulfiram could serve as an efficient copper delivery drug to the brain in Menkes disease, which is characterized by dysfunction in copper transporting (42). Brain images from our study (Fig.1) showed a significant amount of copper accumulated in the brain, which is consistent with the reported results (42).

Recent large clinical data have clearly demonstrated that BAT is tightly linked to obesity and diabetes (4-8). However, imaging methods that can reliably reflect BAT mass are still in shortage. Our data suggested that both ^64^Cu-Dis and [18F]-F-DPA could be used to reliably report on BAT mass, regardless of the activation status of BAT. This is very different from most used BAT tracers such as [18F]-FDG and others, including MRI and optical imaging probes. Interestingly, compared to [18F]-FDG imaging, the images obtained using our method more reliably contour the unique triangle shape of BAT.

Our studies indicate that the BAT contrast is due to the binding of ^64^Cu-Dis to TSPO, which is consistent with previous reports (35, 36). Katz and Gavish reported that Disulfiram and DDC were competitive inhibitors for TSPO or PBR (peripheral benzodianzepine receptor) (35). They demonstrated that Disulfiram and DDC could effectively replace binding of classical TSPO ligands such as Ro5-4864. Gilman et al showed by PET imaging that Disulfiram could reduce the uptake of TSPO ligand 11C-Flumazenil in human brain (36). In spite of the fact that TSPO has been an imaging target for a long time, particularly for brain imaging, very few reports have emerged utilizing it for peripheral target imaging, and biological evaluation of the high TSPO expression level in BAT is rare (41). In the present study, for the first time, we validated TSPO as a biomarker for imaging BAT.

“Browning”, a process of turning WAT into BAT (43-45) represents an exciting approach for converting “bad fat” to “good fat” (BAT and beige fat). Since the “browning” process could result in more mitochondria, and our method is capable of detecting the abundance of TSPO in mitochondria, it is reasonable to speculate that our methods have the potential to monitor the browning process, in which the abundance of TSPO increases with the increase of BAT-like cells.

Although TSPO was discovered nearly 40 years ago, its functions in obesity and adipocytes are not well explored. Our studies discovered and confirmed high TSPO expression in BAT, which opened a new avenue for basic BAT research aimed at investigation of biological functions of TSPO. In addition, since TSPO is tightly associated with inflammation (28), it is conceivable that ^64^Cu-Dis is also suitable for imaging inflammation in different diseases.

In summary, we demonstrated that a synthesis-free method for PET imaging was feasible, and the combination of ^64^CuCl_2_ and Disulfiram could be used for BAT imaging. We also validated, for the first time, TSPO as an imaging biomarker for BAT. We believe that our method can be widely applied for TSPO and BAT imaging, and that the synthesis-free strategy could significantly reduce the cost of PET imaging.

## Materials and Methods

All of the chemicals were purchased from commercial vendors and used without further purification. Disulfiram was purchased from USP (Cat. No. 1224008, Rockville, MD), and dissolved in a solution of 15% DMSO and 85% Cremophor EL (10mg/ml). F-DPA and [18F]-F-DPA were synthesized in Dr. Steven H. Liang laboratory (MGH). Balb/c mice and B6C3F1/J mice were purchased from Jackson Laboratory. All animal experiments were approved by the Institutional Animal Use and Care Committee (IACUC) at Massachusetts General Hospital.

### MicroPET imaging of mice with ^64^Cu-Dis

In vivo PET imaging was conducted with 7-month old B6C3F1/J mice. Mice were anesthetized with isoflurane/oxygen for 5 minutes, and then injected intraperitoneally (i.p.) with 100μL (40 mg/kg) solution of Disulfiram (10 mg/ml). After 60 minutes, mice were injected intravenously (i.v.) with 100 μL ^64^CuCl_2_ (100-150μCi) solution in PBS, pH7.4. After 0.5- and 6-hours, PET imaging was conducted using a 30-minute static imaging on a Sophie Biosciences microPET G4 scanner (Culver City, CA, USA). Imaging analysis was conducted using Amide, a Medical Imaging Data Examiner.

### Cold exposure treatment

Mice (B6C3F1/J) were placed into cold room (4°C) for 2 hours, and then i.p. injected with 100 μL (40 mg/kg) solution of Disulfiram (10 mg/ml). The injected mice were then returned to the cold room for 1 hour, and then i.v. injected with 100 μL ^64^CuCl_2_ (100-150μCi) in PBS, pH7.4. After 30 minutes in the cold room, mice were imaged using a 30-minute static imaging on the G4 scanner.

### Ex vivo bio-distribution of ^64^Cu-Dis

Four-month old Balb/c mice were anesthetized with isoflurane/oxygen for 5 minutes, and then i.p. injected with 100 μL (40 mg/kg) solution of Disulfiram (10 mg/ml). After 60 minutes, mice were i.v. injected with 100 μL of ^64^CuCl_2_ (100-150μCi) solution in PBS, pH7.4. After six hours, mice were sacrificed, and interscapular BAT, interscapular WAT, inguinal WAT, and gondal WAT were excised and subjected to counting in a scintillation counter (Packard Cobra II Auto Gamma Scintillation Well Counter).

### Long anesthesia treatment

Mice (B6C3F1/J) were anesthetized in an induction chamber for 20 minutes, and then i.p. injected with 100 μL (40 mg/kg) solution of Disulfiram. The injected mice were returned to the induction chamber for 1 hour, and then i.v. injected with 100 μL of ^64^CuCl_2_ (100-150μCi) in PBS. After 30 minutes in the chamber, mice were imaged using a 30-minute static imaging on the G4 scanner.

### Blocking studies with TSPO ligand DPA-F

Mice (B6C3F1/J) were i.v. injected with 100 μL solution (3 mg/kg) of F-DPA in 15% DMSO, 15% Cremophor EL and 70% PBS, and then i.p. injected with 100 μL (40 mg/kg) solution of Disulfiram. The injected mice were returned to the cage for 1 hour, and then i.v. injected with 100 μL of ^64^CuCl_2_ (100-150 μCi) in PBS. After 0.5- and 6-hours, mice were imaged using a 30-minute static imaging on the G4 scanner.

### Semi-Quantitative PCR Analysis

Total RNA was extracted from the adipose tissue using TRIzol (Invitrogen, Grand Island, NY). The cDNA was synthesized by iScript^TM^ system (Bio-Rad, Hercules, CA). Reverse transcription reaction was performed using a PTC-100 programmable thermal controller (MJ Research Inc., Waltham, MA). A real time PCR was performed using an iTaq^TM^ universal STBR® green Supermix (Bio-Rad, Hercules, CA) in an Mx3005P qPCR thermocycler (Agilent Technologies, Santa Clara, CA). All values were normalized to cyclophilin expression and further analyzed using the ΔΔ*C*_*T*_ method. The sequences of primers used in the semi-quantitative PCR were as following: TSPO (forward: 5’-CCATGGGGTATGGCTCCTACA-3’, reverse: 5’-CCAAGGGAACCATAGCGTCC-3’); cyclophilin (forward: 5’-GGAGATGGCACAGGAGGAA-3’, reverse: 5’-GCCCGTAGTGCTTCAGCTT-3’).

### Western blotting with anti-TSPO antibody

Adipose tissue was homogenized in a RIPA lysis buffer (Millipore) supplemented with protease and phosphatase inhibitor cocktails (Thermo Scientific). Homogenates were centrifuged at 13,000 r.p.m. at 4°C for 20 minutes, and the supernatants were used as tissue lysates. Protein concentration was determined by BCA protein assay. Seventy micrograms of protein lysate was separated in a 4-20% SDS–PAGE gel. The separated proteins were transferred to a polyvinylidene difluoride (PVDF) membrane, which was then blocked in 5% NFDM at room temperature for 2 hours. After blocking, the membrane was incubated with anti-TSPO primary antibody in TBST buffer (1:1000 dilution, Cell Signaling) at 4°C overnight. After washing with TBST buffer, the membrane was incubated with the secondary antibody (goat anti-rabbit IgG (H+L), Invitrogen) for 1 hour at room temperature. Western Pico Chemiluminescent Substrate (Thermo scientific) was used to visualize the bands. The images were acquired with IVIS®Spectrum (Perkin Elmer, Hopkinton MA) using bioluminescence imaging setting. PageRular Plus Prestained Protein Ladder (Thermo Fisher) (10-250KD) was used as a molecular weight marker.

### Immunohistochemistry with anti-TSPO antibody and histochemical staining with MitoTracker

Adipose tissue was fixed in 4% formaldehyde at room temperature for 24 hours, embedded in paraffin and cut into 5-micron slices. For immunostaining, heat mediated antigen retrieval was first performed using Antigen Retrieval Citra Plus Solution (Biogenex), and then the sections were blocked with 5% NFDM for 1 hour at room temperature. After that, the samples were incubated with anti-PBR/TSPO antibody (Abcam, ab109497) diluted in a blocking buffer overnight at 4°C. After washing with TBST, the samples were incubated with secondary FITC-labeled antibody for 1 hour at room temperature and counterstained with DAPI. For Mitochondria staining, deparaffinized sections were incubated with MitoTracker probes (500 nM) (Invitrogen, MitoTracker® Deep Red FM, cat. No. M22426) for 30 minutes at room temperature. Images were captured using a Nikon ECLIPSE 50i fluorescence microscope.

### In vivo PET imaging with [^18^F]-F-DPA

In vivo PET imaging was conducted with C57BL/6J mice. For cold treatment, mice were placed in the cooled room (4°C) for 2 hours. Mice were i.v. injected with 100 μL of 20-30 μCi [^18^F]-F-DPA in 10% ethanol saline. PET imaging was conducted using a 20-minute static scan on a Sophie Biosciences microPET G4 scanner under short anesthesia (isoflurane/O_2_) (2-3 minutes). Imaging analysis was conducted with Amide.

## Acknowledgement

The authors would also like to thank Pamela Pantazopoulos, B.S. for proofreading this manuscript. We also thank China Scholarship Council of Ministry of Education of China for supporting (J.Y. and J.Y.).

## Supplemental Figures

**SI Fig.1.**
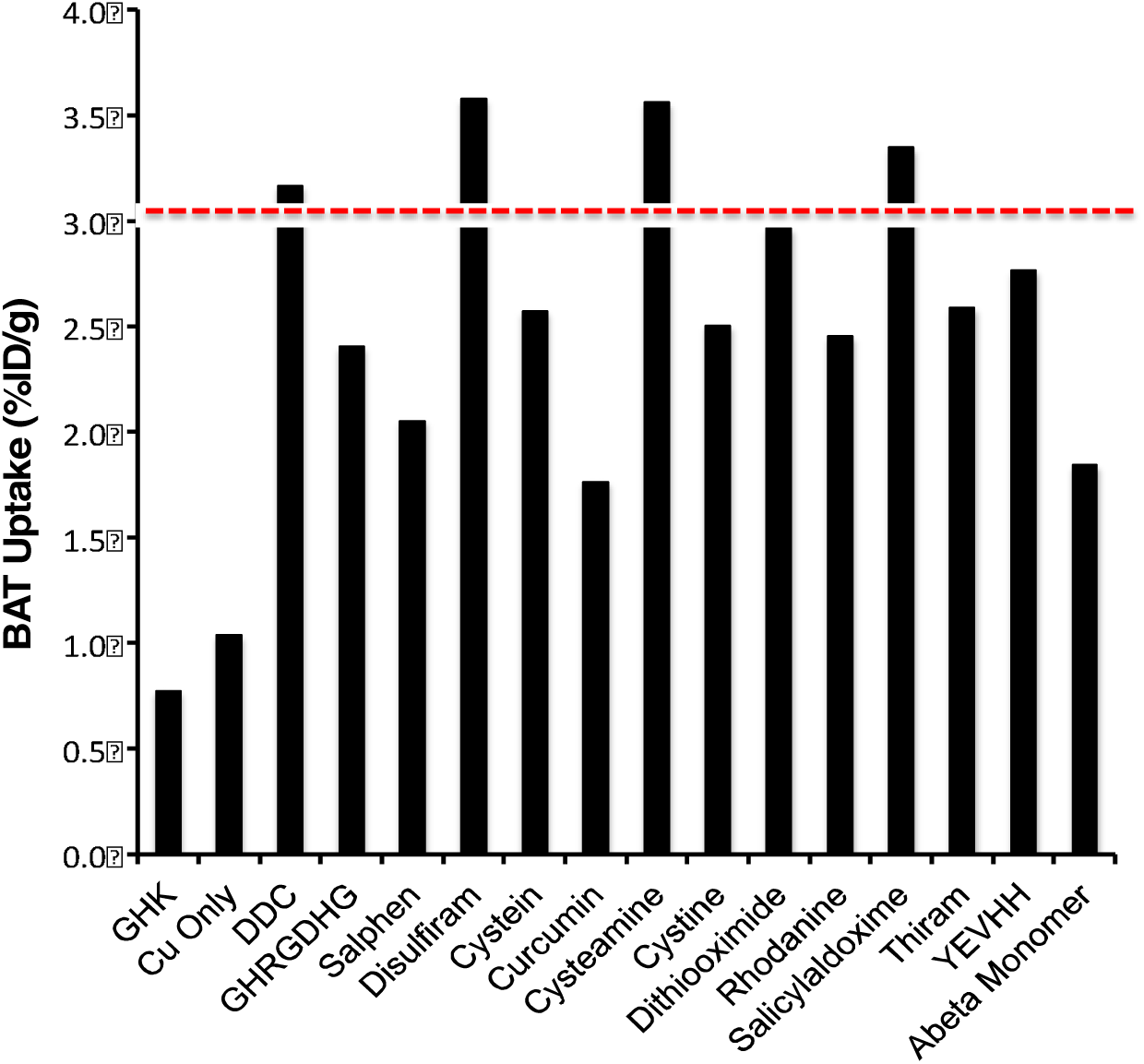
Top-down screening of Copper (II) ligands for BAT imaging.

**SI Fig.2.**
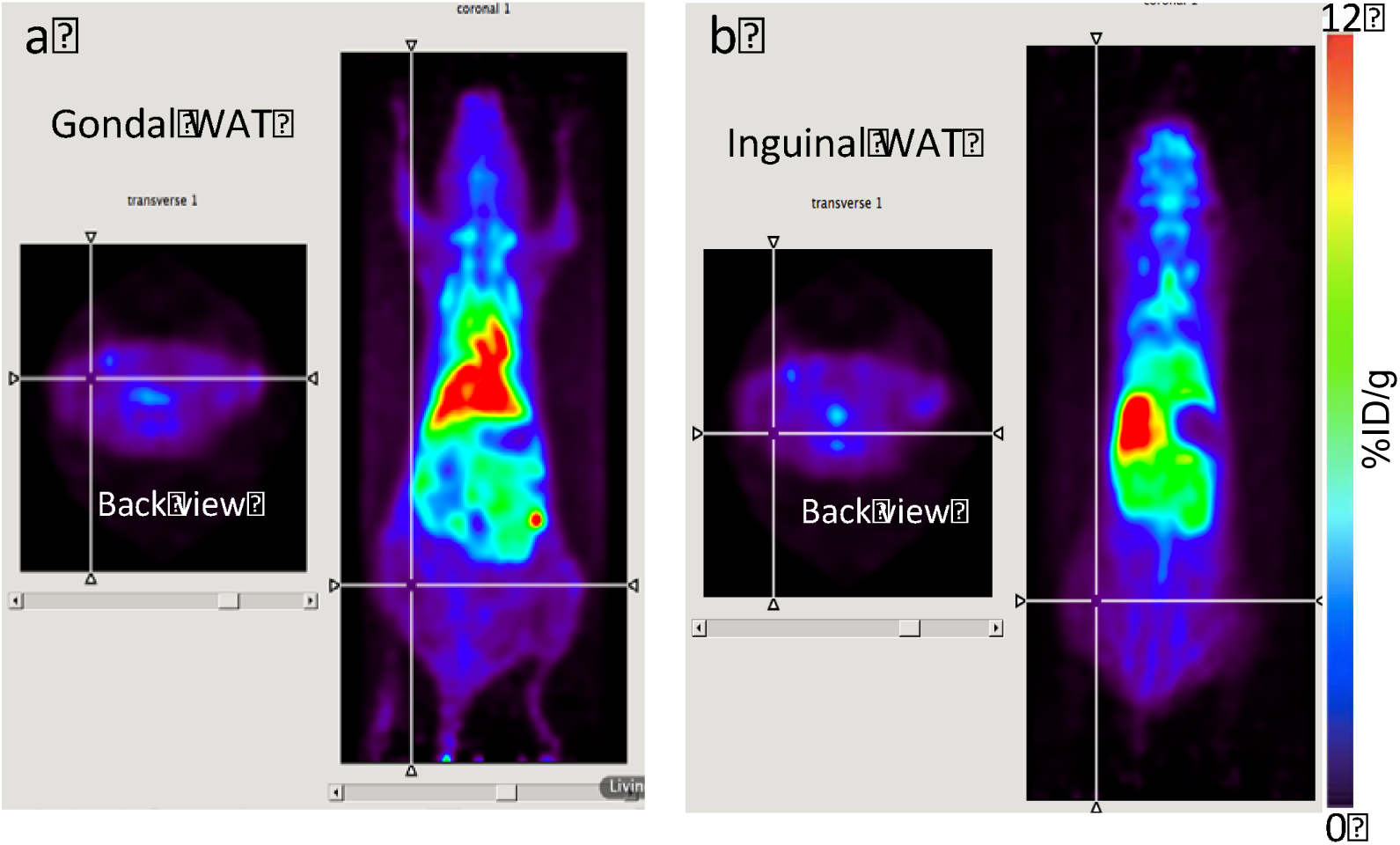
a-b) Representative images of ^64^Cu-Dis uptake in gonadal and inguinal WAT.

**SI Fig.3.**
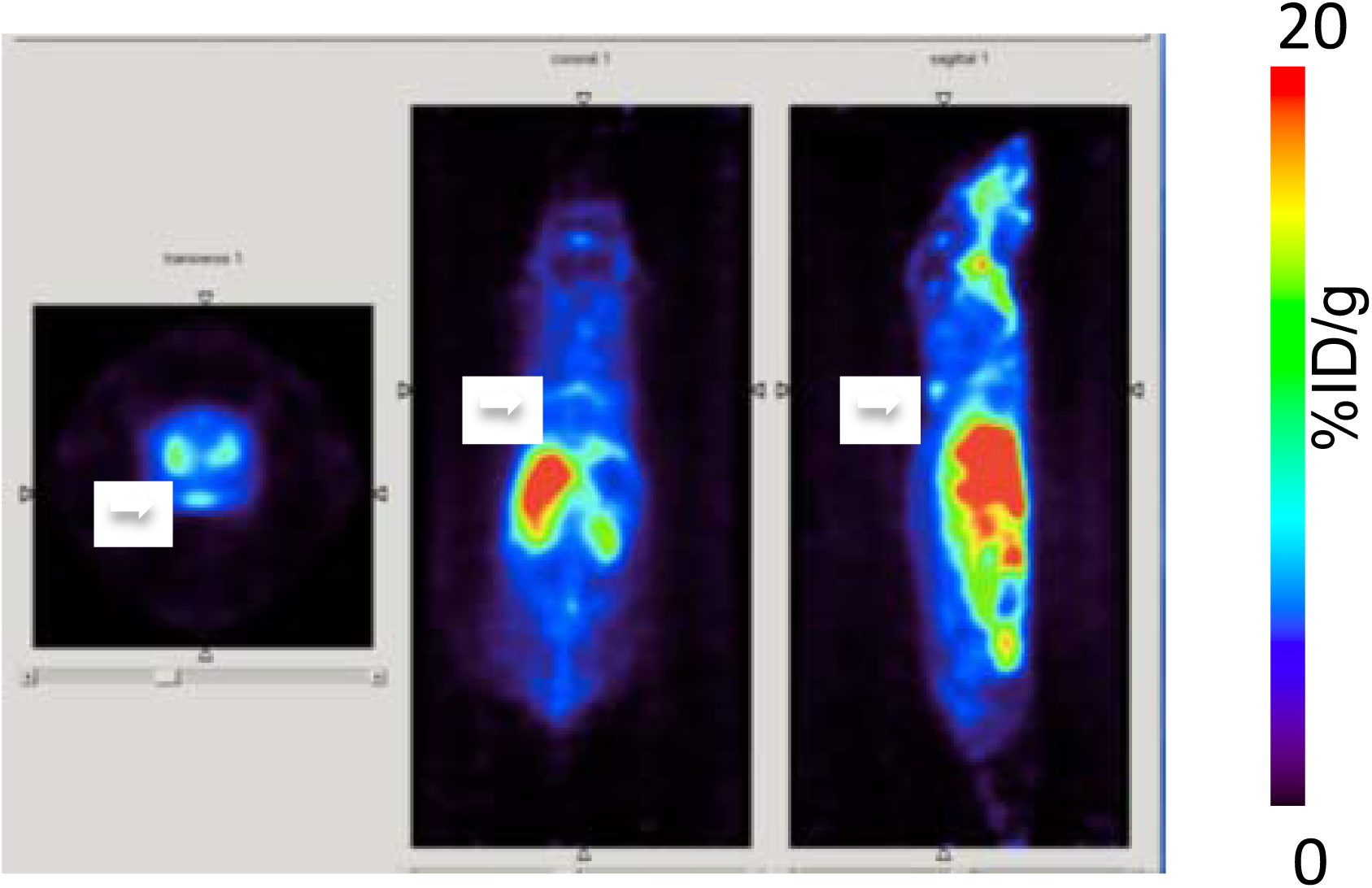
Representative BAT images with ^64^CuCl_2_ only 1 hour after i.v. injection.

**SI Fig.4.**
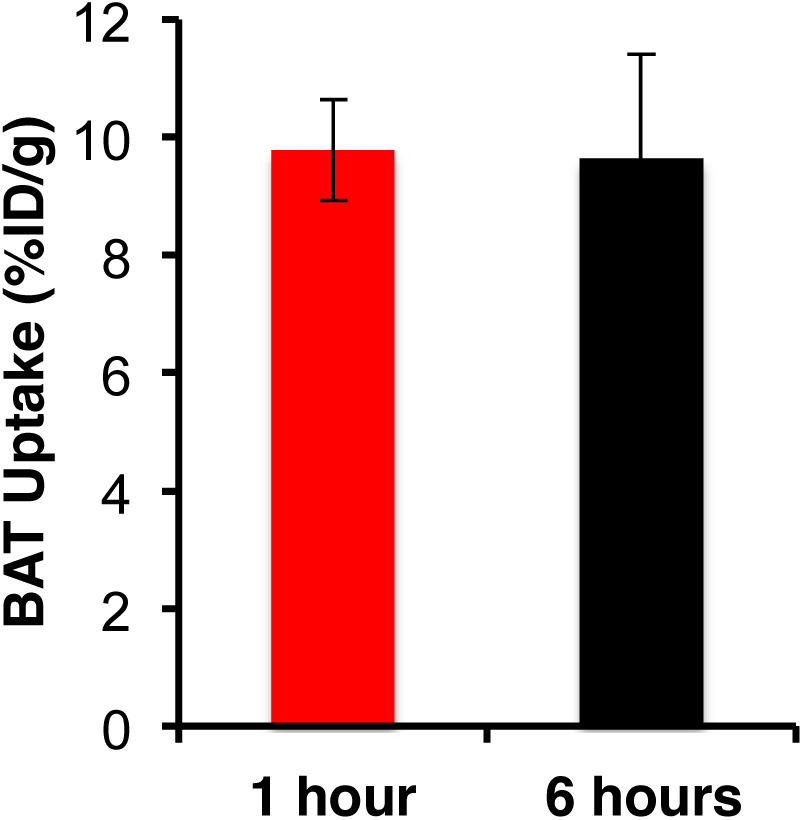
BAT uptake at 1 hour and 6 hours post ^64^Cu-Dis injection.

**SI Fig.5.**
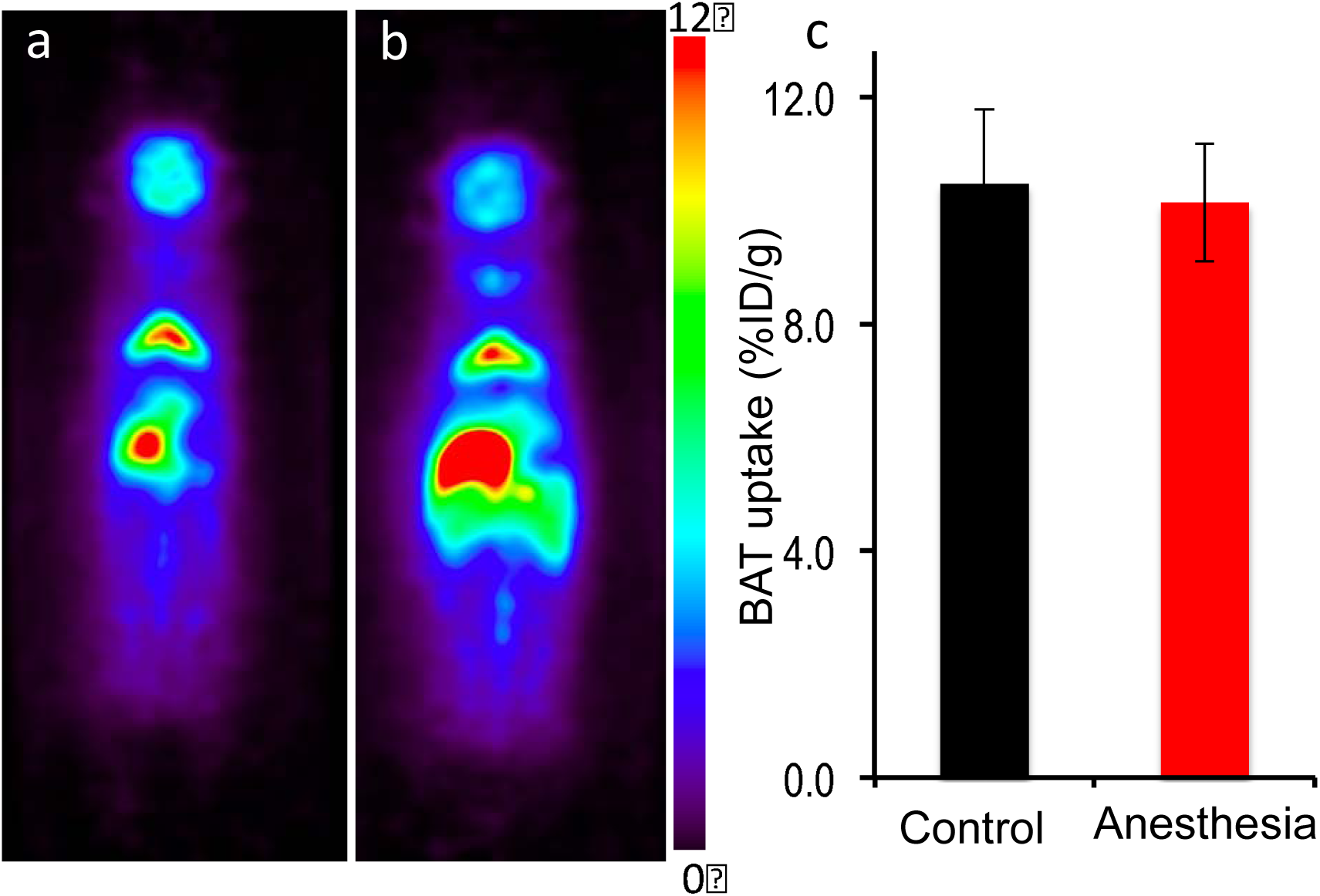
SI Fig.5 Representative BAT images and quantitative analysis of BAT uptake under normal and long anesthesia exposure. There was no significant difference between these two groups.

**SI Fig.6.**
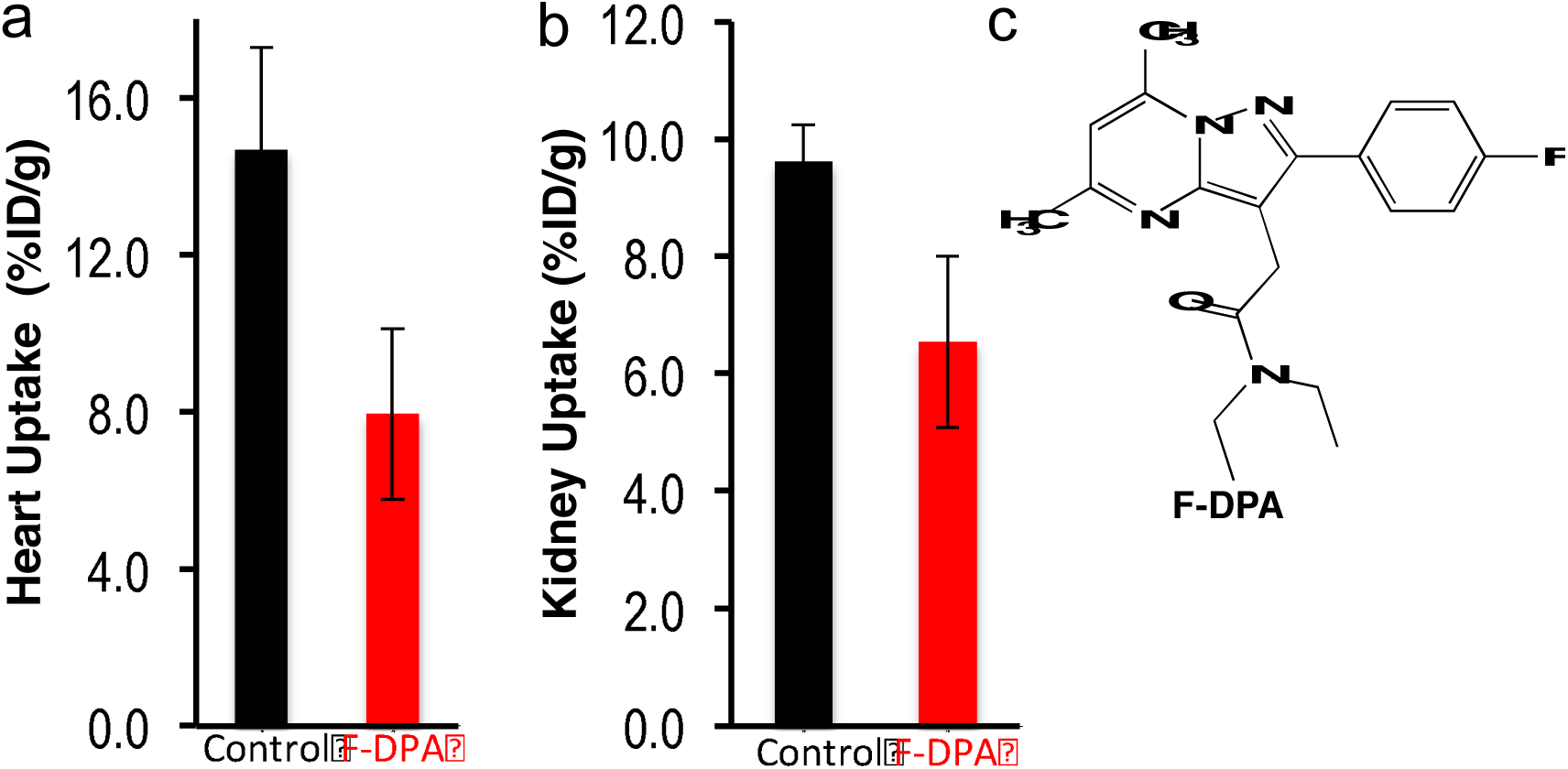
Quantitative analysis of heart (a) and kidney (b) images with F-DPA blocking at 1-hour post ^64^CuCl_2_ injection. (c) Chemical structure of F-DPA.

**SI Fig.7.**
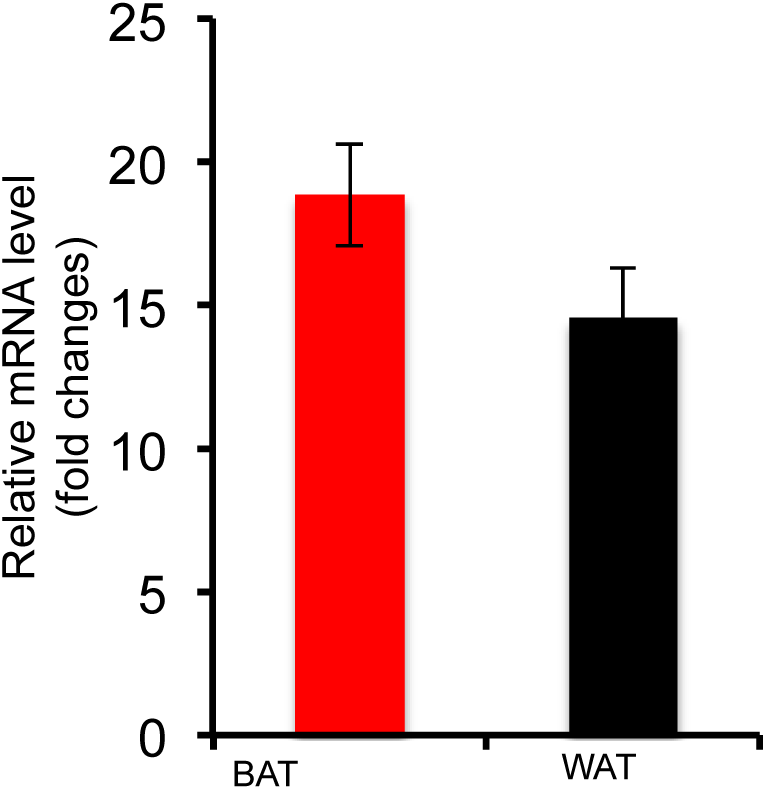
Quantitative PCR analysis of TSPO mRNA level in BAT and WAT tissues.

**SI Fig.8.**
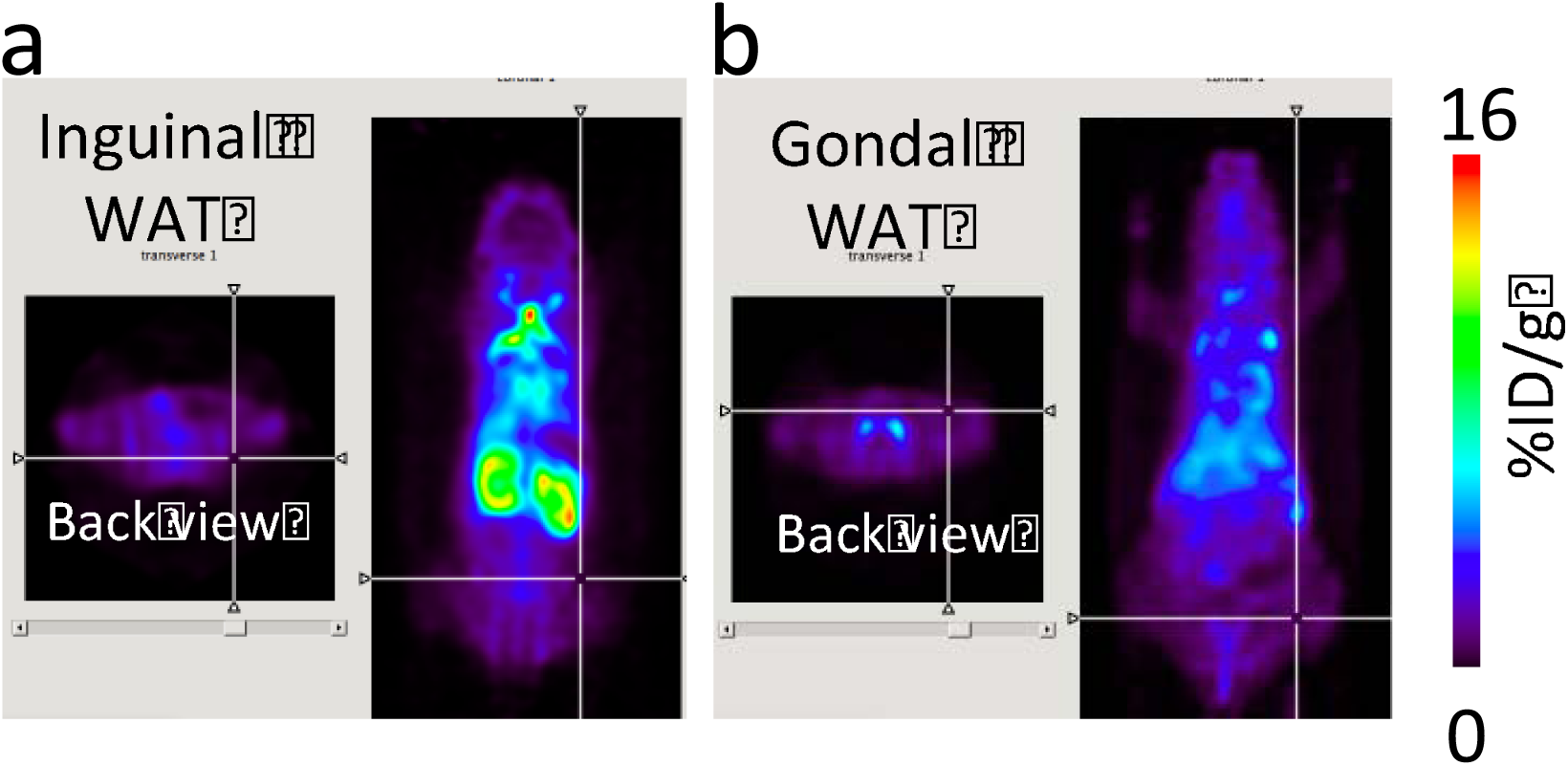
Representative images of 18F-DPA uptake in gondal and inguinal WAT.

**SI Fig.9.**
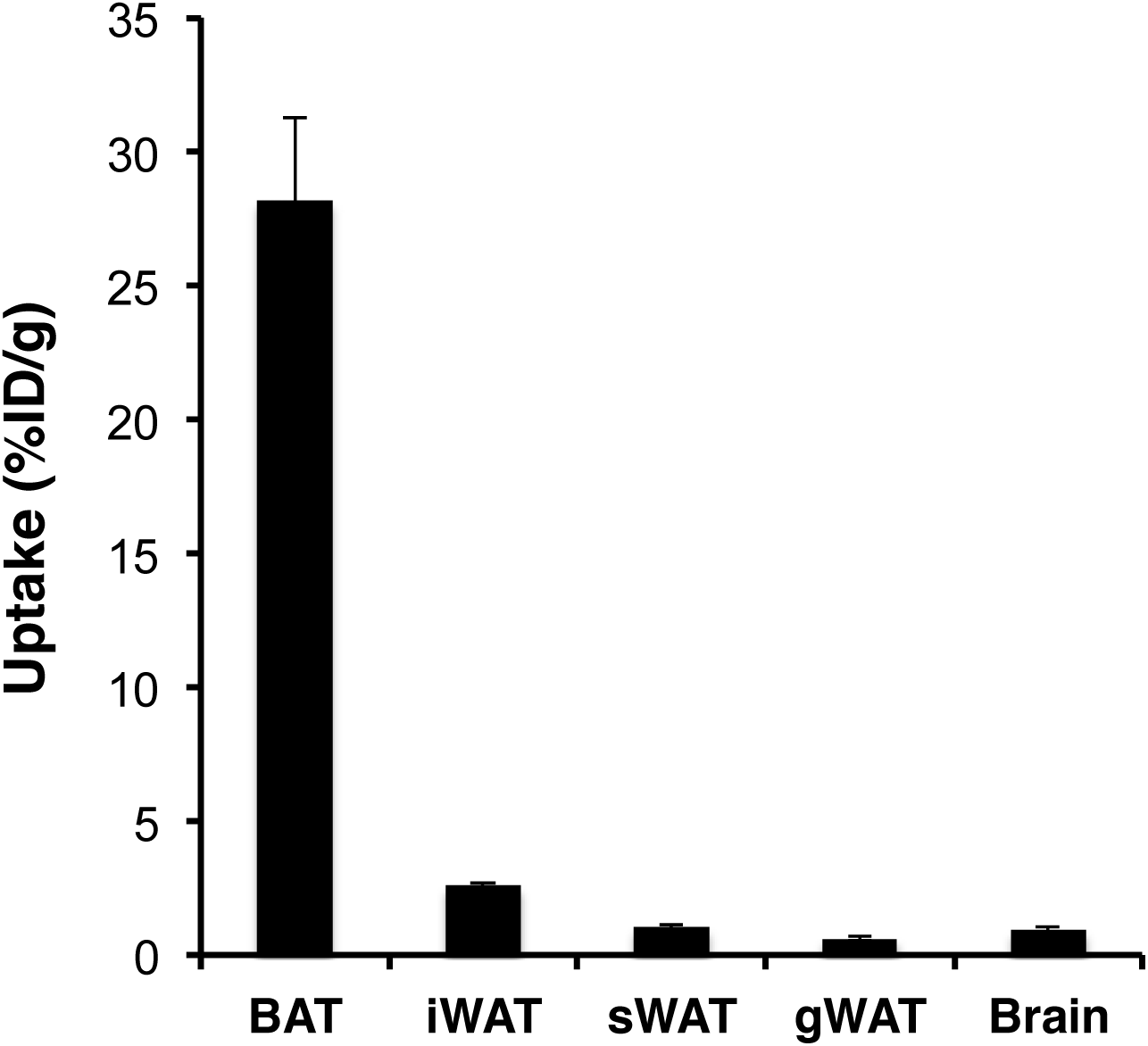
Ex vivo bio-distribution analysis of 18F-DPA uptake in BAT and WAT.

